# Early zygotic engineering promotes targeted large transgene integration and direct production of fully transgenic animals

**DOI:** 10.1101/2024.09.29.615605

**Authors:** Khusali Gupta, Ping Xu, Judith Gallant, Yeonsoo Yoon, Jaime A. Rivera-Pérez, Jeanne B. Lawrence

**Affiliations:** Department of Neurology, University of Massachusetts Chan Medical School, Worcester, MA, USA; Department of Radiology, University of Massachusetts Chan Medical School, Worcester, MA, USA; Department of Pediatrics, University of Massachusetts Chan Medical School, Worcester, MA, USA; Laboratory Animal Sciences Program, Frederick National Laboratory for Cancer Research, Frederick, MD, USA

## Abstract

Genetic engineering has become increasingly feasible with the advent of site-specific nucleases. This technology is highly effective for the generation of small mutations, however targeted insertion of large sequences to generate transgenic animals remains challenging, time consuming, and laborious. Following several failed attempts using the same reagents, we identified a protocol that achieved very high targeted integration of a 3.2 kb transgene into a single locus of a humanized mouse model of Down syndrome. Moreover, we demonstrate in multiple ways that this procedure directly generates numerous non-mosaic founder animals, which bred true to generate all transgenic progeny. *In vitro* fertilization of oocytes was followed by AAV-mediated delivery of donor sequence and electroporation of Cas9 ribonucleoprotein, all parameters designed to promote <u>e</u>arly <u>z</u>ygotic <u>e</u>ngineering (EZE). This strategy can obviate the need to breed mosaic animals for germline transmission and can enable engineering of difficult to breed animals. While efforts to improve genome engineering focus on reagents and delivery techniques, findings here suggest a narrow window of time in the early fertilized oocyte, likely before genome replication, is key to achieve both high integration efficiency and one-step generation of non-mosaic, engineered mice.

## Introduction

For many purposes in biomedicine the ability to insert DNA sequences into the genome is desirable in order to generate transgenic animals. A common practice has been random transgene integration into the genome. This methodology, however, is inefficient and unreliable (Palmiter et al. 1982)). Advances in embryonic stem cells allowed targeted gene insertions using homologous recombination, yet this approach is time consuming and often unsuccessful (Castrop 2010). The development of site-specific nucleases has dramatically improved the efficiency of genome editing, vastly improving small sequence alterations, such as indels or insertions of a few base pairs (H. Wang et al. 2013; Meyer et al. 2010). However, the delivery and targeted insertion of larger DNA fragments, particularly over 1kb has remained a challenge, with the donor fragment size being a critical factor that limits efficacy (Erwood and Gu 2020) Small insertions and deletions are most frequently generated during double-strand break repair by non-homologous end joining (NHEJ), an error prone process active in most cells, including non-cycling cells. In contrast, error-free or precise targeted insertions or deletions using a donor template occurs during homology directed repair, a process generally more limited to proliferative cells (Yeh, Richardson, and Corn 2019; J. Y. Wang and Doudna 2023).

Previously, efforts to generate animals carrying targeted insertions have had to begin with mouse embryonic stem cells (mESCs), since this is technically more challenging in fertilized embryos (Brinster et al. 1989) which have low homologous recombination efficiency (Skarnes 2015). Targeted knock-in of large sequences in mESCs became more achievable with the advent of CRISPR/Cas9 technology, however, this requires a much more prolonged process beginning with ES cell gene targeting, screening for correctly targeted cells, characterization of cell clones, ES cell injection into blastocysts (or morula aggregation), and eventually generation of chimeras. These chimeric animals must then be bred (often for extended periods) to achieve germline transmission to obtain fully transgenic founder animals. This whole process can take up to a year or longer with very low efficiency and without guarantee of getting germline transmission (Carstea, Pirity, and Dinnyes 2009). Moreover, this process is not feasible (or much less successful) for engineering in mouse models that impact early development or fertility, or are more challenging to breed, as is the case for the TcMAC21 mouse model engineered in this study.

Using site-specific nuclease technologies, progress has been made in genetic engineering of mouse zygotes generated by errors in non-homologous end joining to create indels, or for very small insertions with plasmid or linearized DNA (Meyer et al. 2010; H. Wang et al. 2013; Hui Yang et al. 2013); however, targeting of larger transgenes has remained challenging. Engineering in zygotes has generally involved pronuclear injection into naturally fertilized embryos (Meyer et al. 2010; Hui Yang et al. 2013; Gu, Posfai, and Rossant 2018; Abe et al. 2020; H. Wang et al. 2013), a process requiring special expertise and dexterous handling of individual zygotes. An alternative is electroporation-based techniques, which has gained attention given the relative ease in delivering the CRISPR reagents to zygotes. This method has had good efficiency with linearized ssDNA or dsDNA ((Quadros et al. 2017; Hashimoto and Takemoto 2015; S. Chen et al. 2016), however, large kilobase sized transgene knock-in has still been a limitation for getting it delivered into the cell.

More recently, recombinant AAV was shown to facilitate delivery of reagents for genetic engineering in naturally fertilized embryos (Yoon et al. 2018; Chen et al. 2019; Mizuno et al. 2018). AAV delivery avoids the need to remove the zona pellucida from zygotes and produced higher rates of knock-out or knock-in mice. Numerous studies have introduced editing reagents into zygotes and succeeded in producing mosaic mice, which then are bred for germline transmission to generate non-mosaic animals that will “breed true”. Since mosaic animals are routinely produced, this indicates that, despite introduction of all reagents at the single-cell zygote stage, the integration (or editing) event occurred after DNA replication or the first zygotic division. It would, therefore, be an important advance if a procedure could provide not only high targeting efficiency but generate fully edited mice in one-step, avoiding the necessity to breed, especially for animal models with fertility problems.

The findings described here were generated over the course of several efforts to target a 3.2 kb transgene into a unique chromosomal locus in a Down syndrome (DS) mouse model. We sought to target a “minigene” of the long noncoding RNA, XIST, into a human chr21 in the humanized TcMAC21 trisomic mouse, for research on DS (unrelated to genetic engineering techniques). Initially, several protocols were tested, without success, in attempts to target the identical transgene into the same locus (and most using the same gRNA and CRISPR/Cas9 reagents). However, we ultimately identified a protocol that generated exceptionally strong results, providing very high efficacy (66%) of targeted transgene integration, even though only a single target locus (on human Chr21) was present. Furthermore, in our first attempt using the successful protocol we discovered that a remarkably high proportion (9/12) of the transgenic animals were fully transgenic, with no detectable cellular mosaicism. These animals then “bred true” and produced all fully transgenic offspring, rapidly providing multiple founder lines for our research goals (the subject of another study). Although we did not need more mice for our initial purpose, we felt it important to test if we could reproduce similar results using the same protocol, which we did. We detail our methodology here because we believe these findings, and the conceptual considerations they suggest, can be of significant value to others working to utilize or to improve genetic engineering in early embryos.

Over multiple years we attempted to target XIST transgenes into an intron of the DYRK1A gene on human Chr21 in a DS mouse model. This site was targeted successfully in human iPS cells numerous times (using much larger and smaller XIST transgenes)((Jiang et al. 2013; Valledor et al. 2022). In this case there is only one allele of the target locus, which would likely reduce the probability of obtaining a targeted integration event in any given embryo. Here, we begin with briefly summarizing procedures that were not successful to provide perspective on the protocol modifications that ultimately generated surprisingly robust results. All procedures and results described here attempted to target the same transgene into the same locus, hence differences in results are not due to any unusual property of our targeting site or transgene sequence; it was specifics of the approach that made a big difference.

### Attempts using ZFNs and AAV donor delivery in naturally fertilized embryos

We began by attempting to target the 3.2kb donor plasmid by homology directed repair (HDR) using the same zinc finger nucleases (ZFNs) that we previously used successfully to target a ∼14kb XIST cDNA into human iPSCs (and HT1080 cells) *in vitro* (Jiang et al. 2013). Given the success of generating other transgenic mice by AAV infection of naturally fertilized zygotes (Yoon et al. 2018), we packaged our donor plasmid (with homology arms of 502bp and 684bp) and the ZFNs into two separate AAV vectors. We chose the AAV6 serotype as it can transduce pre-implantation embryos well (Yoon et al. 2018).

A small-scale experiment was performed where naturally fertilized zygotes were harvested at 0.5 days post coitum from the TcMAC21 mice, infected with the rAAV vectors, and then implanted into pseudo pregnant females (Figure 1a). As shown in Table 1, 3/11 pups born were GFP+, which is reliably expressed from the human chromosome 21 and thus distinguishes trisomic (Down syndrome) versus wild-type mice in each litter. When these GFP+ pups were screened for transgene insertions, none were found positive. Upon further screening by T7E1 assay to look for small editing by products or indels, we noted 0% indels in the three GFP+ mice.

**Table 1:** Summary of previous attempts targeting the same locus and same transgene using naturally fertilized zygotes for HDR shows little to no efficiency.

| Trial # | No: of zygotes | Delivery Method | Reagents | No: of zygotes transferred | No: of pups born | No: of trisomic (GFP+) pups/ blastocysts* | No: of pups with Indels | No: of pups with KI |
| --- | --- | --- | --- | --- | --- | --- | --- | --- |
| 1 | 39 | rAAV (Donor + ZFN) | ZFN | 34 | 11(+1 dead) | 3 | 0 | 0 |
| 2 | 32 | rAAV (ZFN alone) | ZFN | n/a | n/a | 13* | 0 | 0 |
| 3 | 84 | Pronuclear Microinjection | Cas9 mRNA+ Cas9 RNP+ linearized dsDNA donor | 63 | 30 | 9 | 1(98% -1 del) | 0 |

**Figure 1:**
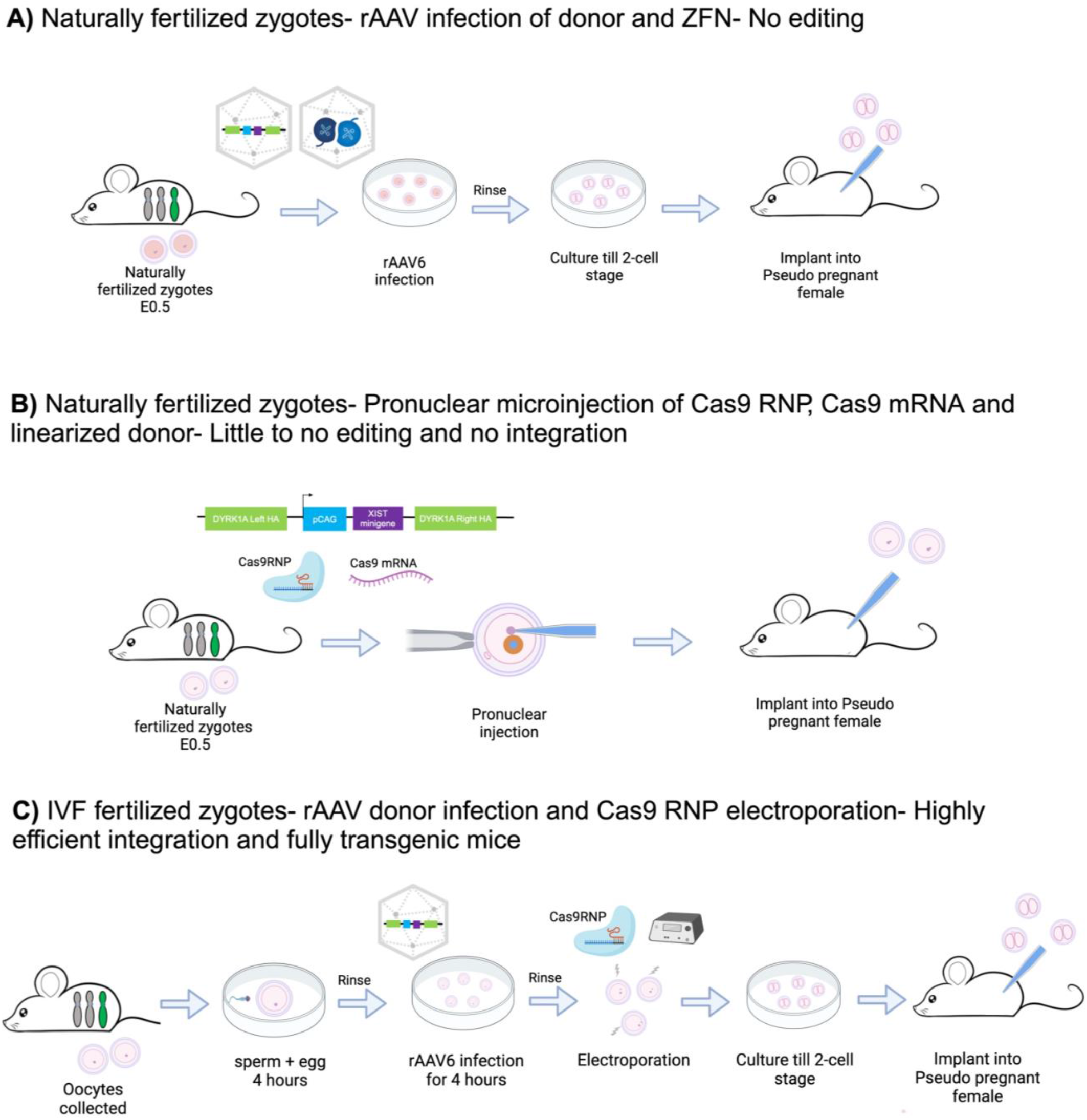
Schematic of different approaches tried to create transgenic mice with targeted integration of a 3.2kb insert at the same locus (DYRK1A intron). Created with Biorender.com **A**) Naturally fertilized zygotes from TcMAC21 females were exposed to rAAV6-DYRK1A-ZFNs and rAAV6-XIST minigene donor, cultured till 2-cell stage and then implanted into pseudopregnant females. **B**) Pronuclei of naturally fertilized zygotes were injected with Cas9RNP and Cas9 mRNA, followed by implantation into pseudopregnant females. **C**) Early zygotic engineering (EZE) in *in vitro* fertilized TcMAC21 zygotes using rAAV donor infection followed by Cas9RNP electroporation. The 2-cell stage embryos were then implanted into pseudopregnant females.

While not detailed here, we had previously attempted to use these same ZFNs to target the large XIST transgene into the same locus in ES cells of the Tc1 DS mouse model, or ZFNs to target a different locus in Ts65Dn mESCs (data not shown). Despite continued success using these same ZFNs in human iPSCs, numerous earlier attempts to use ZFNs in mESCs suggested that ZFNs may be less effective in mouse ESCs (even when the target sequence was identical). Hence, to further test the efficacy of ZFNs in the TcMAC21 mouse line, we performed a blastocyst outgrowth experiment in which naturally fertilized zygotes were treated with rAAV transduction of the ZFNs alone and cultured *ex-vivo* for blastocyst outgrowths. As summarized in Table 1, we did not find any editing by T7E1 assay.

In view of the lack of success of ZFNs in mice (both *in vitro* and *ex vivo)*, we decided to switch to the cheaper and more easily programmable CRISPR nucleases.

### Pronuclear Microinjection of CRISPR/Cas9 (RNP+ mRNA) +Linear donor into naturally fertilized embryos

As several studies have generated transgenic animals using pronuclear microinjection of Cas9 components (Abe et al. 2020; Gu, Posfai, and Rossant 2018), we identified gRNAs for the same target sequence (as above). Cas9 (in both RNP and mRNA forms) along with our linearized dsDNA donor, were injected together into the female pronucleus and cytoplasm of each individual naturally fertilized embryo (Figure 1b). As shown in the Table 1, 9/30 pups born were GFP+, but none had our transgene inserted. However, we did have one GFP+ pup that upon screening showed ∼90% indels, encouraging us to continue with CRISPR/Cas9 rather than ZFNs.

However, since this rate of editing was not high, to increase chances of generating at least one transgenic TcMAC21 mouse, we decided to avoid pronuclear microinjection, which requires time consuming and laborious manipulation of each embryo. Instead, we improvised to try rAAV donor delivery coupled with electroporation of SpCas9 protein and sgRNA (in RNP form) which allows several embryos to be treated at one time, rather than individually manipulated. Use of these rAAV (Yoon et al. 2018) or rAAV and electroporation (Chen et al. 2019) for reagent delivery into naturally fertilized zygotes was shown to substantially improve editing frequency to generate mosaic mice, which could then be bred to obtain germline transmission for non-mosaic founder mice. As will become clearer below, a significant distinction from our earlier attempts and these prior studies is that for our third attempt we used oocytes that had just been fertilized *in vitro*, rather than more typical naturally fertilized zygotes flushed from the oviduct.

### Highly Successful Targeted Integration in IVF Embryos using AAV donor delivery and Cas9 RNP electroporation

Given limited availability of female mice (as TcMAC21 males are sterile), for our next attempt we maximized use of small numbers of females by introducing reagents into *in vitro* fertilized zygotes instead of naturally fertilized zygotes. As shown in Figure 1c, 4 hours post fertilization (sperm and eggs were setup), the fertilized zygotes were exposed to rAAV donor for 4 hours followed by RNP electroporation. Results were significantly more successful even though we were targeting the same sequence with the same reagents, as shown in the Table 2. At this very first try combining rAAV donor transduction and electroporation of Cas9 RNP into IVF embryos, of 18 trisomic mice born, PCR showed that 12 had targeted insertion of the large (3.2kb) transgene at the desired site (Figure 2b). In 66.67% of mice PCR amplification of the 5’ and 3’ ends showed clear transgene insertion at the target site (Fig 2b). Three independent founder lines were Sanger sequenced using PCR to examine both sides of the integration site and this showed integration precisely in the same target sequence, without edits to the flanking sequences (Fig 2c). It is noteworthy that there is only a single target locus on the one human chromosome, which would reduce (possibly by half) the chance of hitting the desired locus if two chromosomes or alleles were present. Hence, results show ∼67% targeting efficiency of all alleles, whereas most studies report frequencies for either of two alleles. (By extrapolation, results suggest that the chance of inserting into both alleles, if present, would be ∼45%.). We had one GFP+ mice that had partial transgene insertion at the target site and another GFP+ mice that could be an off-target insertion (detected only donor integration but no homology).

**Table 2:** Targeting *in vitro* fertilized zygotes generates high efficiency of transgenic mice.

| No: of zygotes AAV treated | No: of two-cell embryos transferred | No: of pups born | No: of trisomic (GFP+) pups | No: of pups with KI (%) | No: of KI pups with CRISPR cut site detected (mosaic) | No: of KI pups with no CRISPR cut site (non-mosaic) |
| --- | --- | --- | --- | --- | --- | --- |
| 161 | 143 | 32 | 19* | 12 (66.67) | 3 | 9 |

**Figure 2:**
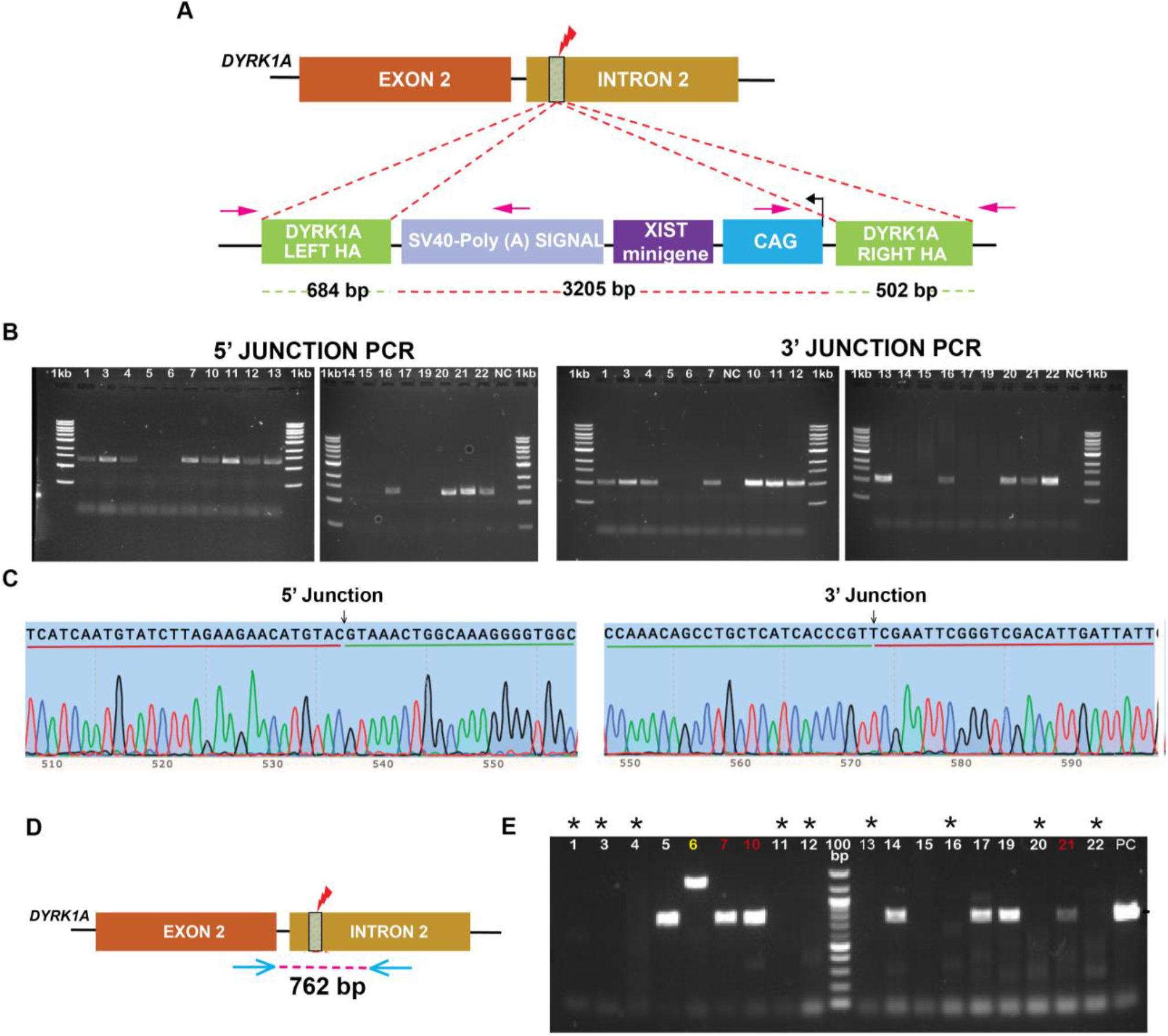
Analysis of targeted integration in IVF zygotes shows high integration efficiency and revealed generation of fully transgenic non-mosaic mice. **A**) Schematic of transgene targeting the intron of human DYRK1A locus (on the single human chr21). The HDR donor is a 3.2kb insert sequence that contains the CAG promoter running the XIST minigene and a SV40-PolyA sequence cassette flanked by left homology arms of 684bp and right homology arms of 502bp. HA, homology arm and pink arrows represent primers flanking the junction regions. **B**) PCR genotyping confirms correctly inserted 5’ and 3’ junctions of the modified DYRK1A locus in 12/18 GFP+ (trisomic) mice generated by EZE (Fig 1c). 5’ junction expected band size 1314bp and the 3’ junction expected band size 1026bp. NC, negative control; 1kb, 1kb marker ladder. **C**) Representative Sanger sequencing and chromatograms for correctly edited mice, n=3. The green underline marks the left or right homology sequence, and the red underline marks the insert sequence on the trace. **D**) Schematic of strategy to detect gene editing (indels) or WT at the DYRK1A target site (Pink lightning arrow). Blue arrows represent primers targeting the flanking CRISPR cut site. **E**) PCR genotyping of the DYRK1A CRISPR cut site (as shown in Fig2d) identifies WT or edited loci (NHEJ) that did not receive the knock-in. The * marks samples that were negative for the CRISPR locus showing no WT allele indicates generation of non-mosaic transgenic mice in 9/12 KI mice. The red labeled samples indicate the mice that are mosaic as they are positive for the cut site as well as for the targeted KI (Fig2B). We also identified sample #6 in yellow that had a greater than expected product size which upon examination had partial insertion of the donor.

### One-step production of fully-transgenic founder mice directly from IVF Embryos

The presence of only one human chr21 allele in the TcMAC21 mouse also facilitated evaluating the extent of mosaicism in the mice, since any given cell could have just one type of allele. While PCR (in Table 2) was used to show the transgene was inserted into the target locus in 12 mice (two-thirds), we used PCR primers flanking the target site to detect cells with an intact target site (no transgene integration) in each individual animal (Fig 2d). It was initially surprising that in 75% (9/12) of the knock-in mice, PCR to the uninterrupted target region could not be detected (Fig 2e). Tail tip DNA was used for this analysis, as it represents many cells, including potentially different developmental lineages (skin, bone, cartilage, blood vessels, nervous tissue). It was unexpected that the procedure would generate in “one-step” fully transgenic (non-mosaic) animals, and at a very high rate, and for a large 3.2kb transgene insertion. Hence, we took further steps to investigate and validate this possibility.

An important validation came when we bred 6 out of our 9 knock-in (KI) females, which showed that *all* trisomic off-spring (born at the expected Mendelian ratio) also carried the targeted 3.2kb insertion (Table 3). As shown in Table 3, *all* mice that appeared to be fully transgenic by PCR produced pups (45 in total) that were also fully transgenic, and founders have bred true for several generations. We conclude that the protocol not only provided high efficiency for targeted integration of a 3.2kb transgene, but also provided the large advantage of producing fully transgenic founders in a single step from the trisomic IVF embryos. This one experiment generated nine fully transgenic founder mice, which can provide several major advantages, as considered in the Discussion.

**Table 3:**
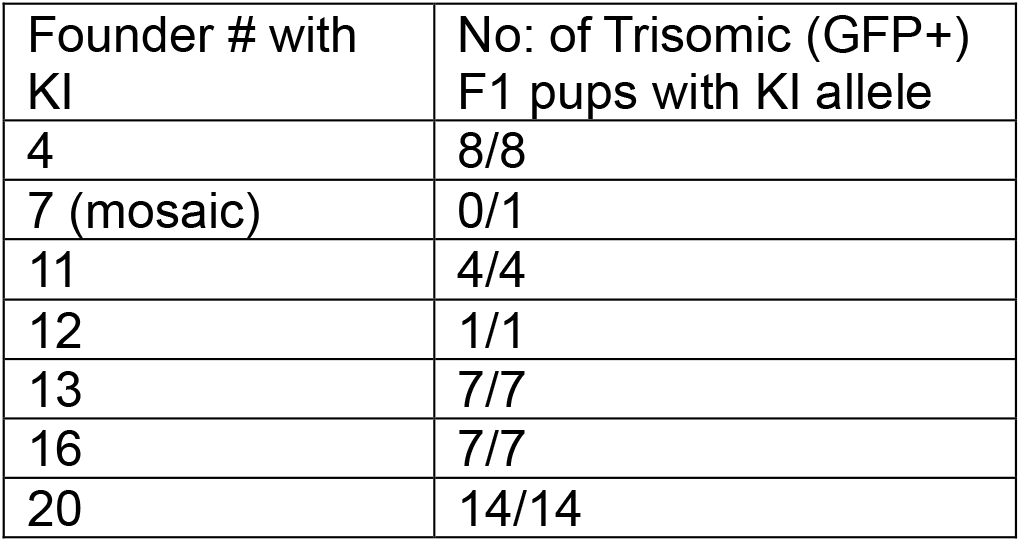
Analysis of founder female breeding and genotype of F1 mice with Knock-In.

### Reproducible high efficiency and generation of fully transgenic mice in whole blastocyst embryos

The high frequency of fully transgenic mice indicated to us that our introduction of donor and CRISPR reagents into the IVF embryos very likely resulted in transgene integration occurring at the one-cell stage, before the first cell division. Since the efficient generation of fully transgenic mice was unanticipated, especially after several failed attempts with the same transgene, target site, and CRISPR reagents, we tested if we could reproduce similar results. Since we surmised timing of integration at the one cell stage was critical, we again used *in vitro* fertilized zygotes, but this time cultured them to blastocyst out growths, to screen both embryonic and extra-embryonic components for cells with or without the transgene insertion.

In the first experiment (above), the rAAV donor was introduced to all oocytes four hours after exposure to sperm. Since the actual sperm penetration and post-fertilization steps and maturation can vary by a few hours, AAV donor was introduced to what would be a mixture of oocytes in earlier and later stages of fertilization and post-fertilization changes. In the second experiments we were more focused on the importance of precise timing, so we tried to more precisely define and control it, by using polar body formation to indicate which oocytes had been fertilized and begun post-fertilization steps. Hence, at four hours AAV donor was introduced to oocytes that had reached polar body stage (Group I), and for the less advanced oocytes that had not yet formed polar bodies (Group 2) the time before AAV donor infection was extended to 8 hours. By bracketing the timing in this way we hoped not to miss a critical time window and potentially also provide insights into a potentially narrow window of zygote/pronuclear state that allowed highly efficient targeted integration.

Importantly, as summarized in Table 4, this experiment demonstrated that the protocol was able to reproduce similar highly efficient results for targeted insertion of the 3.2 kb transgene. Approximately 65% of all trisomic blastocysts carried the transgene at the target site in both groups I and II. Results further affirm that this process can generate fully transgenic and non-mosaic animals. Collectively this second experiment produced a major fraction (37%) of knock-in mice that were fully transgenic (7 of 19). We note that in the first experiment an even higher fraction (75%) of knock-in mice were fully transgenic (12/18), which, might reflect slight modifications in timing and oocyte stage, as will be considered in the Discussion.

**Table 4:** Summary of targeted transgene integration in timed *in vitro* fertilized zygotes and analysis at blastocyst stage. Based on visual observations oocytes with extrusion of polar body and beginning formation of prominent pronuclei were selected as group I and transduced with AAV four hours after sperm exposure. The rest of the oocytes (Group II) that had not yet formed polar bodies were exposed to AAV donor eight hours later. In all cases Cas9 RNP electroporation was done four hours post-AAV infection.

| Groups | Trisomic (GFP+) blastocyst | No: of blastocysts with KI (%) | No: of mosaic blastocysts | No: of non-mosaic blastocyst |
| --- | --- | --- | --- | --- |
| Group I (4hr pf) | 12 | 8 (67) | 5 | 3 |
| Group II (8hr pf) | 17 | 11 (64) | 7 | 4 |

Collectively, these results demonstrate that the protocol for early zygotic engineering (EZE) detailed here can generate both efficient integration efficiency of a large transgene and moreover, generate fully transgenic mice in one-step method, without the need to breed mosaic animals to generate full founder mice.

### Analysis of small allelic edits and mosaicism in founder derived mice

The above analyses demonstrate that this early stage of the zygote generates very high efficiency of transgene integration which occurs by HDR typically during S-phase. Many genetic engineering efforts seek to induce smaller changes for gene “knock-outs”, generated by the error-prone process of NHEJ (not requiring S-phase). The cell state of the early zygote (approximately Pronuclear PN1-PN3 stage, Figure 3a) may be particularly receptive to HDR because of constituents within the oocyte cytoplasm as discussed below. To gain further insights, we examined mice from the first experiment using Inference of CRISPR Edits (ICE) analysis to detect small edits at the target site in mice that were either mosaic or had no transgene. We also assessed how many different types of alleles were detected in individual mice, and the relative proportions of allele types, giving further insight into the stage when editing occurred (before or after cell divisions). In the 25% (3/12) of mosaic pups (i.e., some cells with KI and some with WT allele by PCR) it is likely that integration occurred at the two-cell stage or after DNA replication. When we further examined by ICE (Table 5), these three mosaic KI mice all had just one type of other allele (either an indel or unedited allele). Results suggest that these mosaic animals had a 50:50 split, consistent with the editing events happening either at the 2-cell stage or in G2 of one-cell stage. Extending this approach to all the rest of the GFP+ trisomic pups (which lack transgene knock-in), we found the editing efficiency detected by ICE was extremely high, with 8 out of remaining 9 mice having small edits (insertions/deletions) at the target site, as indicated in the table 5. This shows that this “early zygotic stage” is extremely receptive not only to transgene integration by HDR, but for small edits introduced by NHEJ. Results also suggest edits occurred early, because we did not detect more than 2 different alleles in any individual GFP+ pup, and 3 of 5 mice with indels were essentially non-mosaic (99-100% of cells with same edit), and only one mouse had a high number of different alleles. Mouse #14 had a 3:1 ratio (74% -7bp del: 24% WT), consistent with edits occurring at the two cell-stage or late G2 phase (with skewing due to either asymmetrical division or correction of one sister chromatid as template). These results again emphasize the ability to achieve non mosaic founders by targeting zygotes soon after fertilization.

**Table 5:** Summary of all target site edits and allelic proportions.

| GFP+ Mice ID # | Editing byproducts by Inference of CRISPR Edits (ICE) analysis | Allelic variations detected | Types of Alleles | Editing outcome | Timing of editing relative to DNA replication | Potential timing of when editing occurred |
| --- | --- | --- | --- | --- | --- | --- |
| 1 | n/a | KI | One | Non-mosaic | Pre-replication | One-cell |
| 3 | n/a | KI | One | Non-mosaic | Pre-replication | One-cell |
| 4 | n/a | KI | One | Non-mosaic | Pre-replication | One-cell |
| 5 | -7bp del (100%) | -7bp | One | Non-mosaic | Pre-replication | One-cell |
| 6 | Lots of indels | Partial insertion | Several (>12) | Mosaic | Post-replication | After two-cell G2 |
| 7 | +1 ins (99%) | i) KI<br>ii) +1bp | Two | Mosaic | Post-replication | One-cell G2 or early 2-cell |
| 10 | WT (98%) | i) KI<br>ii) WT | Two | Mosaic | Post-replication | One-cell G2 or early 2-cell |
| 11 | n/a | KI | One | Non-mosaic | Pre-replication | One-cell |
| 12 | n/a | KI | One | Non-mosaic | Pre-replication | One-cell |
| 13 | n/a | KI | One | Non-mosaic | Pre-replication | One-cell |
| 14 | -7bp del (74%)<br>WT (24%) | i) -7bp<br>ii) WT | Two | Mosaic | Post-replication | One-cell G2 or 2-cell |
| 15 | none | none | n/a | none | n/a | n/a |
| 16 | n/a | KI | One | Non-mosaic | Pre-replication | One-cell |
| 17 | +1 ins (98%);<br>-1 del (2%) | i) +1 bp<br>ii) -1 bp | Two | Mosaic | Post-replication | One-cell G2 or early 2-cell |
| 19 | -1 del (100%) | -1 bp | One | Non-mosaic | Pre-replication | One-cell |
| 20 | n/a | KI | One | Non-mosaic | Pre-replication | One-cell |
| 21 | WT (99%) | i) KI<br>ii) WT | Two | Mosaic | Post-replication | One-cell G2 or early 2-cell |
| 22 | n/a | KI | One | Non-mosaic | Pre-replication | One-cell |

**Figure 3:**
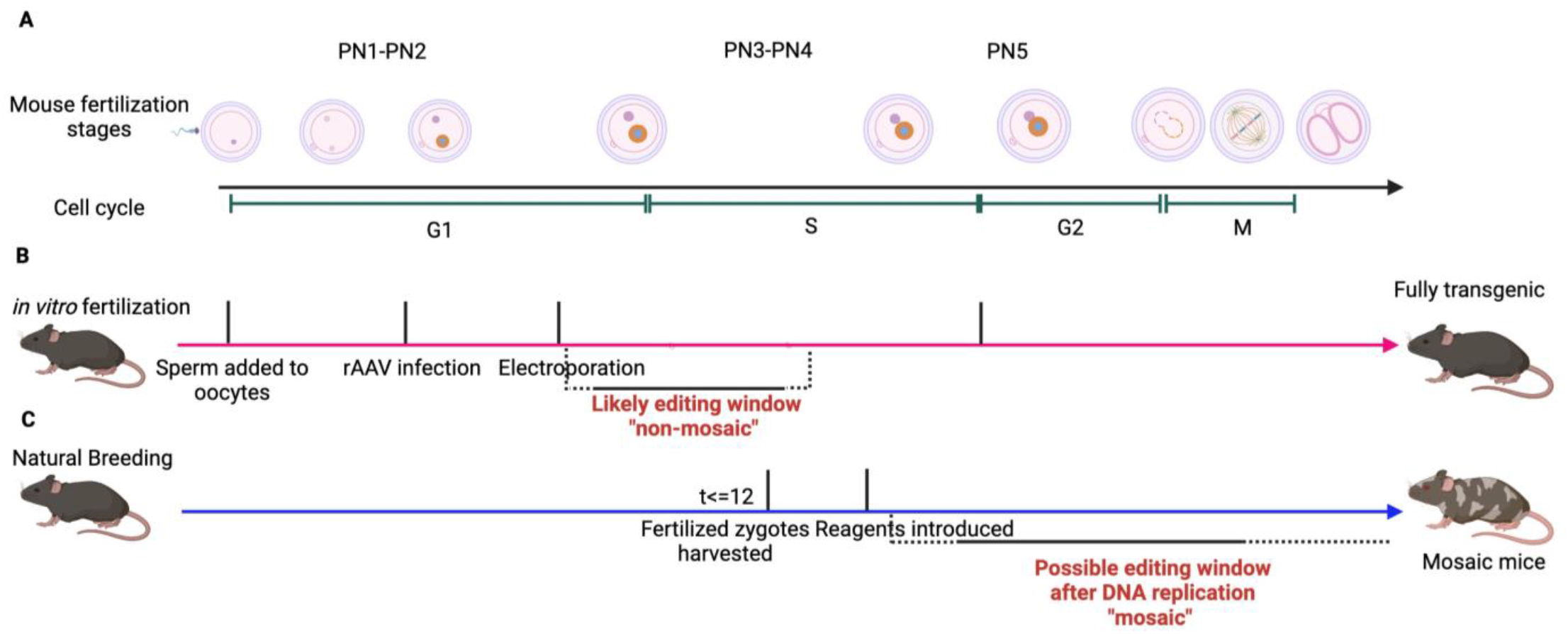
Mouse post-fertilization stages and approximate time windows for integration in IVF vs naturally fertilized zygotes to generate non-mosaic or mosaic animals. Created with Biorender.com **A**) The visible representative stages of mouse fertilization after sperm meets the egg. Following sperm head penetration into the oocyte, the male and female pronuclei begin to appear faintly at the periphery (PN1-PN2). They slowly become more prominent and begin to move towards each other. The two pronuclei replicate their DNA separately as they move closer (PN3-PN4), and then fuse at the G2 phase (PN5). Subsequently, the nuclear envelope breaks down followed by mitosis and generation of two-cell stage embryos. This process takes approximately 22 hours in mice. **B**) Using *in vitro* fertilization, early pronuclear zygotes at PN1-PN2 are exposed to rAAV donor transduction followed by electroporation 4 hours later. All reagents are delivered within several hours of fertilization (G1 phase) and are ready to act. The likely window of when KI occurs to produce non-mosaic fully transgenic mice is in G1/early-S prior to replication of the target locus. **C**) Naturally fertilized zygotes are more advanced with their DNA replicated, based on when the mating occurs. Thus, the window of early zygotic editing is missed and mostly mosaic mice are generated even if rapidly acting reagents are used.

In summary, all these results reinforce that this modified protocol provides an extremely high rate of editing as well as low or no mosaicism. While the Cas9, gRNA, and target sequence were the same in our failed attempts, dramatically more successful results were obtained with a procedure that used *in vitro* fertilized embryos instead of naturally fertilized embryos. The difference in timing post-fertilization between IVF and naturally fertilized zygotes is about 8 to 9 hours, which is actually a large developmental difference relative to the rapid progression of changes during early stages following fertilization. Figure 3A outlines the known biology of stepwise changes immediately following sperm penetration into the oocyte, and Fig 3B and 3C shows the approximate timing difference between natural versus *in vitro* fertilized oocytes. These steps include dramatic remodeling and decondensation of both sperm and oocyte chromatin, formation of male and female pronuclei, and progressive movement of pronuclei closer together until they ultimately fuse to form the single nucleus of the zygote. Importantly, DNA replication occurs separately in the male and female pronuclei *before* they fuse, and by the time naturally fertilized zygotes are harvested the pronuclei have generally already begun replicating their DNA. Hence, the state of the zygote when editing reagents are introduced will differ markedly between IVF versus naturally fertilized embryos. How these timing and staging differences likely contribute to the marked differences in results we obtained is considered in the Discussion.

## Discussion

This study demonstrates a genetic engineering procedure that promotes extremely efficient targeted integration of a large (3.2 kb) transgene, but remarkably also produced numerous *non-mosaic, fully* transgenic mice *directly from fertilized oocytes*. This approach allowed us to rapidly produce numerous mice from multiple founders in a trisomic DS model that is challenging to breed (sterile males). Our goal was accomplished when we generated nine fully transgenic founder mice from one IVF experiment, however, because the approach we identified could be valuable to others, we decided to better document and analyze our methodological findings. The ability to surpass breeding requirements to obtain fully modified animals overcomes a major hurdle generally and makes it more feasible to study mutations that impact fertility or development. Even for mice that are easier to breed, this reduces time, costs, and labor, and can reduce genetic “bottle necks” by generation of multiple independent founders. These issues pose an even greater barrier for modification of larger animals which have longer gestational periods (e.g. rhesus macaques, cows, pigs, etc.). Implications of our findings, as discussed below, have significance for an increasingly large area of genetic engineering in research and agriculture for which improved technologies are sought (McFarlane et al. 2019). While our findings are limited to a particular locus in the DS mouse model but targeting into this locus repeatedly failed by different procedures, and these differences are instructive. Based on the robustness of results we repeatedly obtained, and the logic of the information described below, we believe key principles of our protocol can also provide a framework for improving transgene integration and gene editing in other contexts. (We note these principles have recently been applied by the UMASS Chan Transgene core for highly successful genetic engineering at other loci in a different mouse model.)

### Early timing as Key: Genome editing in *in vitro fertilized* oocytes

Several prior attempts to insert the same transgene into the same locus in naturally fertilized zygotes failed to produce even one mosaic animal (e.g. with some cells carrying the transgene). As described in results, we stepwise modified our protocols to incorporate certain technical aspects from other studies, including use of AAV donor delivery ((Yoon et al. 2018)) and electroporation of Cas9 ((S. Chen et al. 2019)). However, these studies, which used naturally fertilized zygotes, produced mosaic mice. In contrast, the procedure identified here delivered not only highly efficient integration but many fully transgenic mice. Several considerations discussed below support that the central aspect of our protocol which produced these results is due to small but critical differences that impact *the timing of when genetic modifications occur relative to S-phase in the fertilized oocyte or the first cell division*. This, in turn, is determined by three general factors: 1) The rapidly changing maturation of the post-fertilization oocyte, particularly in relation to DNA replication. 2) When reagents are introduced relative to the changing state of the fertilized egg. 3) How quickly the editing reagents can act, and the donor sequence is available, for integration.

Most efforts to create genetically modified animals use naturally fertilized zygotes, however, as outlined in Figure 3, these naturally fertilized zygotes will be substantially more “mature” than IVF oocytes. Thus, when naturally fertilized zygotes are harvested, they are already well into S-phase (and beyond) before editing reagents are even introduced. In contrast, our protocol uses IVF which provides a much longer interval before S-phase, G2, and onset of cell-divisions. The timing of delivery and speed with which reagents act will further define when genetic changes occur relative to the developmental stages of the zygote or early embryo. Our results suggest that the quick availability of our reagents for rapid genome editing also contributed to the optimal timing in the early fertilized oocyte. Use of Cas9 RNP allows for quick action of the enzyme, but also provides a short half-life of Cas9 protein, which avoids any lingering of the Cas9 that can produce more mosaicism. Electroporation of Cas9 is easier to perform quickly on multiple embryos simultaneously and delivers reagents accessible to both pronuclei. Additionally, rAAVs provide efficient delivery of donor templates simultaneously to numerous oocytes, and AAVs have been suggested to facilitate HDR, possibly due to their single stranded nature and potential impact of the inverted terminal repeats (ITRs) (Hirsch 2015). Notably, the AAV donor delivery occurred only four hours after mixing sperm and oocytes, so oocytes were at an early post-fertilization stage.

Clearly, the specific reagents and steps in our protocol are designed to promote genetic changes in the early zygote. However, as explained below the nature of our results further supports our conclusion that early zygotic engineering (EZE) is optimal to generate non-mosaic, fully transgenic animals.

### A narrow window of time before DNA replication is likely optimal to avoid mosaicism

Transgene integration by canonical HDR (homology directed repair) occurs in late S-phase or G2 because this process repairs the double-strand break using homology between sister chromatids (or donor template) in the presence of specific repair enzymes and proteins (Rad51, Ctip etc)(Han Yang et al. 2020). Gu, Posfai, and Rossant 2018 concluded that the long G2 of the two-cell stage was optimal for integration by HDR, however this timing will predictably produce substantial mosaicism. A study by Abe et al. 2020 utilized *in vitro* fertilized zygotes and emphasized the importance of introducing reagents at the S-phase (PN3-PN4) for achieving highly targeted KI. Despite targeting early pronuclear stages, Abe et al did not screen for or discuss mosaicism as they may be hitting a time frame after the DNA had started replicating. When integration by HDR occurs in late S/G2 at the one-cell stage, the transgene integration occurs only in one of the two chromatids, thus producing daughter cells with and without the transgene. Logically, however, fully transgenic mice would arise if the integration event occurred *prior* to DNA replication, so that the single genome that will give rise to all cells of the body is modified.

The fact that 75% of our transgenic mice were fully transgenic indicates that this was not an anecdotal event or a “fluke” but occurred repeatedly (and independently) in 9 of 12 fertilized oocytes. To demonstrate reproducibility, we tested two repetitions analyzed at the blastocyst stage, which produced similarly strong results. Robust generation of fully transgenic animals was proven not only based on PCR in mouse tail tips or entire blastocysts, but by demonstrating that the transgenic mice directly from zygotes then bred true, since all of their many trisomic offspring carried the large insertion. We also showed by sanger sequencing of three independent non-mosaic founders that the intact transgene inserted precisely at the same target site in all three. To our knowledge production of fully modified animals that breed true has not been demonstrated previously. Given overall targeting efficiency of 66%, it is very unlikely that two independent integrations occurring on both sister chromatids in each post-replication zygote 9 out of 12 times. For reasons further explained below, we believe these results indicate an optimal (relatively narrow) window for targeted integration may exist in G1 or early S, prior to replication of the target locus. Consistent with this, in our first experiment which produced the highest fraction of non-mosaic animals more oocytes were at an earlier stage because we added AAV-donor four hours after adding sperm, irrespective of polar body formation. Based on visual observations we estimate reagents were introduced in pronuclear stages PN1-PN3 for the first experiment (see Fig 3b), immediately after the appearance of pronuclei. However, for the second experiment we thought S-phase was key, so we selected fertilized oocytes that had already formed polar bodies and added AAV at 4 hours, and then waited longer (8 hours) for the other fertilized oocytes to mature. These results are consistent with our suggestion that the earlier timing, likely prior to S-phase or replication of the target locus, is optimal for producing fully transgenic animals.

We also did ICE analysis of alleles lacking the knock-in in all the mice produced from our first IVF experiment. This further showed that all mice carried some modification at the target locus and showed that there were only one, or two, types of alleles present in any given mouse. This indicates that even smaller edits are likely occurring at the one cell stage zygote, and some before replication. Most studies do not systematically screen for allelic variations or cellular mosaicism rates since the common and expected outcome/goal is mosaic animals (requiring breeding). Given that there was only one target allele on the human chr 21 in the TcMAC21 mice, this facilitated our analysis of cellular mosaicism, which clearly shows that frequently the genetic change introduced was present in all cells of the animal. Hence, we suggest a protocol that provides for early introduction and action of genetic engineering reagents, especially prior to replication, a key for the generation of non-mosaic founders.

### Unique biology of the fertilized oocyte may provide for non-canonical mechanisms

Homology directed integration, or small edits by NHEJ, will be impacted by the accessibility of chromatin to nucleases, and availability of DNA repair factors. A study by (Gu, Posfai, and Rossant 2018) proposed that G2 of the two-cell stage was optimal for knock-in partly because chromatin would be more accessible during zygotic genome reactivation at this stage. However, both the male and female pronuclei undergo even more dramatic chromatin remodeling and decondensation during early steps following fertilization (Khokhlova et al. 2020). It is the common practice that gene editing using pronuclear injection introduces reagents into the male pronucleus, given that i) it is typically larger and more visible than female pronucleus and 2) chromatin of male pronucleus is repackaged from protamines to histones, thus thought to be more amenable for genome editing. However, results presented here show that the female pronuclear genome is as receptive, if not more, to genetic engineering.

Since in TcMAC21 mice, the human Chr21 is only transmitted through the mother, this is an advantageous system to assess changes that occur in the female pronucleus, reflecting its distinctive chromatin state and capacity for DNA repair. Our results demonstrating CRISPR targeted cuts in 17/18 animals shows that the female pronucleus is highly accessible to nuclease targeting, and the high fraction that integrated the transgene (12/18) shows a very high integration rate. Studies have reported that oocytes contain high concentrations of repair proteins that facilitate DNA repair during and post fertilization, before zygotic genome activation begins. In contrast, sperm have no DNA repair and hence rely on oocytes repair proteins after fertilization (Khokhlova et al. 2020; Stringer et al. 2018). Based on other observations in cells (Schep et al. 2021; E. Chen et al. 2022) heterochromatin marks at the site of DSB promotes HDR, which suggests that the high H3k9trimethylated DNA in transcriptionally inactive female pronuclei (Funaya and Aoki 2017) may contribute to higher HDR accessibility. Additionally, there are processes (such as transposition of mobile elements like LINE-1 that are specifically active during the early stages of zygote development (Jachowicz et al. 2017). All these points highlight the unique nature of genome remodeling in fertilized oocytes, which may involve non-canonical mechanisms that promote homology mediate genome repair.

As HDR is a precise repair pathway widely thought to be restricted to the S and G2 phase of cell cycle when the repair proteins and sister chromatids are available as a template. However, as we introduce our reagents at the G1/S transition phase and our donor template is accessible, we think we can achieve non-canonical HDR as early as the start of S-phase or prior to the replication of the targeted loci. Previous work by Hashimoto, Yamashita, and Takemoto 2016, show the ability to generate non-mosaic mutant embryos with a 3bp restriction site inserted by HDR by targeting IVF zygotes as early as possible post fertilization in G1. In addition, the recent reports of rAAV facilitating HDR even in post-mitotic cells (Bijlani et al. 2021; Ishizu et al. 2017; Nishiyama, Mikuni, and Yasuda 2017) and also the identification of a new homology mediated end joining (HMEJ) repair mechanism active in other cell cycle phases(Owen et al. 2021; Yao et al. 2017), we think non-canonical HDR could occur outside the late S and G2 window in “the special early fertilized oocytes”. Hence, despite knowing the exact repair mechanism involved our study points to a narrow window of time that precedes S phase and is most optimal for generation of non-mosaic transgenic animals.

### Broader implications of non-mosaic genetic engineering in IVF embryos

Mice are small animals which, in most cases, breed rapidly at low cost, however the ability to circumvent mosaicism has importance well beyond mice. As noted above, mosaicism is a major hurdle for engineering in other mammals that are long and expensive to breed. Our study emphasizes use of IVF embryos, because naturally fertilized zygotes will generally be too advanced to capture the optimal early pronuclear stage. Recent work using IVF was able to generate bovine embryos with little to no mosaicism (for indels or targeted KI) utilizing the HMEJ pathway. While most studies are limited to embryos, ours is the first to demonstrate non-mosaic transgenic animals that also bred true, producing all transgenic progeny, over multiple generations.

IVF technology has become increasingly effective and mainstream, led by expanding clinical applications for human assisted reproduction. Several strong ethical considerations regarding gene editing of IVF embryos are currently debated, and concerns about mosaicism is one of the obstacles (Brokowski and Adli 2018; Turocy, Adashi, and Egli 2021). While other important issues remain, the results and concepts presented here provide a basis for designing procedures that circumvent or minimize mosaicism. We suggest that the pre-zygotic stage of the fertilized oocyte (prior to nuclear fusion) not only supports large transgene integration but is also optimal to avoid mosaicism. Hence, the special properties of the fertilized oocyte, coupled with advances in genetic engineering technologies, may ultimately lead to promising new avenues for human therapies.

## Acknowledgements

We would like to thank Dr. Miguel Sena Esteves and his lab for packing and small-scale preparation of our rAAV vectors. The work described here was supported by grants from the National Institute of Health R01HD094788 to J.B.L. and American Heart Association predoctoral fellowship 897828 to K.G.

## Materials and Methods

### Mouse strains

All experiments involving animals were conducted following the institutional animal care and use committee at the University of Massachusetts Chan Medical School. TcMAC21 (RBRC 05796) mice were obtained from RIKEN BRC, Japan; BDF1 males were obtained from Jax labs (Strain #100006), and Swiss webster mice were obtained from Taconic Biosciences. All animals were maintained in a 12-hour light cycle.

### Collection of naturally fertilized zygotes

For naturally fertilized zygotes, 4-12 weeks old TcMAC21 females were super-ovulated by intraperitoneal injection of pregnant mare serum (PMSG) and 47 hours later with human chorion gonadotropin (hCG). Super-ovulated females were mated with BDF1 males at a 1:1 ratio. The middle of the light cycle of the day when a mating plug was observed was considered embryonic day 0.5 (E0.5) of gestation. Zygotes were collected at E0.5 by tearing the ampulla with forceps and incubating in M2 medium containing hyaluronidase to remove cumulus cells.

### *In vitro* fertilized zygotes

For *in vitro* fertilized zygotes, 4-8 weeks old TcMAC21 females were super-ovulated using CARD HyperOva (Cosmo Bio, KYD-010-EX) following the manufacturer’s instructions. Briefly, 0.1-0.2ml of CARD HyperOva is injected intraperitoneally into 4-8 weeks old female mice at 5pm. After 48 hours, 7.5 I.U i.p injection of human chorionic gonadotropin (hCG) is carried out,13 hours post hCG injections, the oocytes are harvested. 6-month-old BDF1 male donors were used for sperm collection.

### Plasmid construction

#### Donor Plasmid CAG-XIST minigene

The CAG promoter was PCR amplified from the CAG-eGFP plasmid from Addgene (Plasmid # 11150) and the XIST minigene, DYRK1A homology arms and vector backbone was adapted by PCR from the pTRE3G-A-repeat-Ef1a-RFP::DYRK1A plasmid (Valledor et al). The five PCR products were GIBSON assembled using the NEBBuilder HiFi DNA assembly kit (NEB, E2621). The vector was then assembled to incorporate the AAV2 ITR sequences from U1a Nme2Cas9 rAAV (Addgene, Plasmid #129534). The primer sequences are listed in the Primer table.

#### Linearized dsDNA donor

For pronuclear microinjection, the CAG-XIST minigene donor was digested using restriction enzymes Not1-HF (NEB, R3189S) and HindIII-HF (NEB, R3104S) to make a linearized dsDNA donor fragment and the DNA was gel extracted using QIAquick Gel extraction kit (Qiagen, 28704).

#### Human DYRK1A ZFN Plasmid

The two human DYRK1A targeting ZFNs used in (Jiang et al. 2013), were merged into one plasmid with a T2A fragment PCR amplified from AAVS1 ZFN plasmid. The plasmid was put into the AAV2 ITR plasmid backbone U1a Nme2Cas9 rAAV (Addgene, Plasmid #129534. Primer sequences are listed in the table.

### Recombinant AAV vectors production

Briefly, recombinant AAV vectors were produced by calcium phosphate triple transfection of plasmids in HEK293 cells. For small scale preparation, approximately 1.7×10^8 cells were transfected and rAAVs were purified by using iodixanol gradient centrifugation (Burger and Nash 2016). Briefly, HEK293 cells were detached by vigorously shaking the culture flask. The cell suspension underwent three cycles of freezing (dry ice/ethanol bath) and thawing (37dC water bath) for cell lysis and subsequent benzonase treatment. After centrifugation, the supernatant was transferred to an ultracentrifugation tube containing a discontinued gradient of 15,25 40 and 60% of iodixanol (Accurate Chemical, Cat No: AN1114542). Gradient centrifugation was carried out at 504,000xg for 70 min at 20dC. The rAAV vectors at the 40-60% interface were collected and subjected to desalting using a Zeba column (Thermo Fisher Scientific, Cat No UFC910024). All rAAV vectors were titrated by droplet digital PCR (ddPCR, Bio-rad) for genomes and silver staining of capsid proteins.

### sgRNA design

sgRNA targeting the human DYRK1a locus were designed using CRISPOR (Concordet and Haeussler 2018) and commercially purchased from IDT. The sequence was 5’-CACCCCTTTGCCAGTTTACA-3’ (2nmol). The PAM sequence is CGG.

### Pronuclear injection of naturally fertilized embryos

Zygotes derived from crosses between TcMAC21 superovulated females and BDF1 males were collected at E0.5 as mentioned above. DYRK1A sgRNA (20ng/ul), the recombinant SpCas9 nuclease (IDT, Cat No 1081058) (50ng/ul) and SpCas9 mRNA Trilink; L-7206 (50ng/ul) were injected into the female pronucleus and cytoplasm along with linearized dsDNA XIST minigene donor (5ng/ul) using the FemtoJet 4i microinjector (Eppendorf). Injected zygotes were transferred into the oviduct of pseudo pregnant Swisswebster females that were plugged on the same day.

### RNP Assembly

To assemble the Cas9/sgRNA RNP complex, the recombinant SpCas9 nuclease (IDT, Cat No 1081058) was incubated in a 1:1.5 molar ratio with sgRNA to obtain a final concentration of 8uM Cas9/sgRNA RNP in a 10ul solution containing 20mM HEPES, pH 7.5 (Sigma, H3375), 150mM KCl (Sigma, P9333), 1mM MgCl2, (Sigma, M4880), 10% glycerol (Sigma, G5516), and 1mM reducing agent tris (2-carboxyethyl) phosphine (TEP, Sigma, C4706). This mixture was incubated at 37dC for 10 minutes before immediate use.

### Recombinant AAV vector transduction

Fertilized zygotes were incubated in 10 or 15ul drops of KSOM (Potassium supplemented Simplex Optimized Medium, Millipore, MR-106) containing the following rAAV vectors: rAAV6.CAG-XIST-minigene (1.24×10^7GCs/ml); rAAV6-DYRK1a-ZFNs (1.24×10^7GCs/ml) for 4 hours. Drops were placed in 35mm plates under mineral oil (Sigma, M8410) at 37dC in a tissue culture incubator containing 5% CO2 and 5% O2. After the incubation the embryos were rinsed once in M2 medium. AAV transduced embryos were either cultured till blastocysts outgrowths, transferred to pseudopregnant females (in the case of ZFNs) or electroporated with the assembled RNP complex (CRISPR-READI)(S. Chen et al. 2019).

### Electroporation of Cas9RNP

Briefly, zygotes were transferred to 40ul of Opti-MEM reduced serum media (Thermo Fisher, 31985062), and 10ul of Opti-MEM containing the embryos were mixed with 10ul assembled SpCas9 protein/sgRNA RNPs. The 20ul embryos and RNP mixture was transferred to a 1-mm electroporation cuvette (Bio-RAD, 1652089) and electroporated (square wave, 6 pulses, 30 V, 3 ms duration, 100 ms interval) using a gene Pulser XCell electroporator (Bio-Rad, 1652660). Zygotes were recovered and transferred to fresh KSOM drops and cultured overnight. Treated zygotes that developed to two-cell stage were either transferred surgically into the oviducts of pseudopregnant Swiss Webster (Taconic Biosciences) females to generate mice or cultured in KSOM for 3 days to form blastocysts or as blastocyst outgrowths. Blastocysts or tail snips from pups were then collected for lysis and DNA prep to perform PCR analysis.

### Genotyping analysis of edited embryos and mice

Crude DNA extract was obtained from blastocyst stage embryos or embryo outgrowths by placing each individual embryo in 10ul of embryo lysis buffer (100ug/ml Proteinase K, 50mM KCL, 10mM Tris-HCL, 2.5mM MgCl2, 0.1mg/ml gelatin, 0.45% v/v IGEPAL and 0.45% v/v Tween 20), followed by lysing at 55dC for 30 mins to 2 hours and heat inactivated at 95dC for 8 mins.

Mouse tail DNA was extracted by proteinase K digestion overnight at 56dC followed by heat inactivation at 95dC for 10 mins, followed by phenol chloroform genomic DNA extraction and purification. Briefly, equal volume of Phenol/chloroform/isoamyl alcohol was added to the lysed DNA sample, mixed and then spun down for mins at max speed. For all PCR reactions, Q5 Hot Start High-Fidelity 2X Master Mix (NEB, M0494S) was used and the PCR was setup in a Bio-Rad Thermocycler.

For screening of donor integration and knock-in experiments, primer pairs as mentioned in the Table below were used for PCR genotyping of the 5’ and 3’ junctions consisting of one primer pair from within the knock-in inserted sequence and the other flanking the homology arm at the DYRK1a locus.

### T7E1 nuclease assay and Indel analysis by Inference of CRISPR Edits (ICE)

The target site of the DYRK1a locus gene was amplified using Q5 Hot Start High-Fidelity 2X Master Mix (NEB, M0494S), purified using the QIAquick PCR Purification Kit (Qiagen, 28106) and subjected to T7E1 nuclease assay according to the manufacturer’s instructions (NEB, M0302L). Digested products were resolved on a 1-2% agarose gel containing SYBR safe (Invitrogen, S33102) and imaged.

To screen for CRISPR induced indels, PCR products flanking the target cut site were purified using the Qiagen QiaQuick PCR purification kit and submitted for Sanger sequencing. The sequencing files were then analyzed using Synthego’s Performance Analysis, ICE Analysis. 2019. v3.0. Synthego; [June 2022]

#### PCR primers

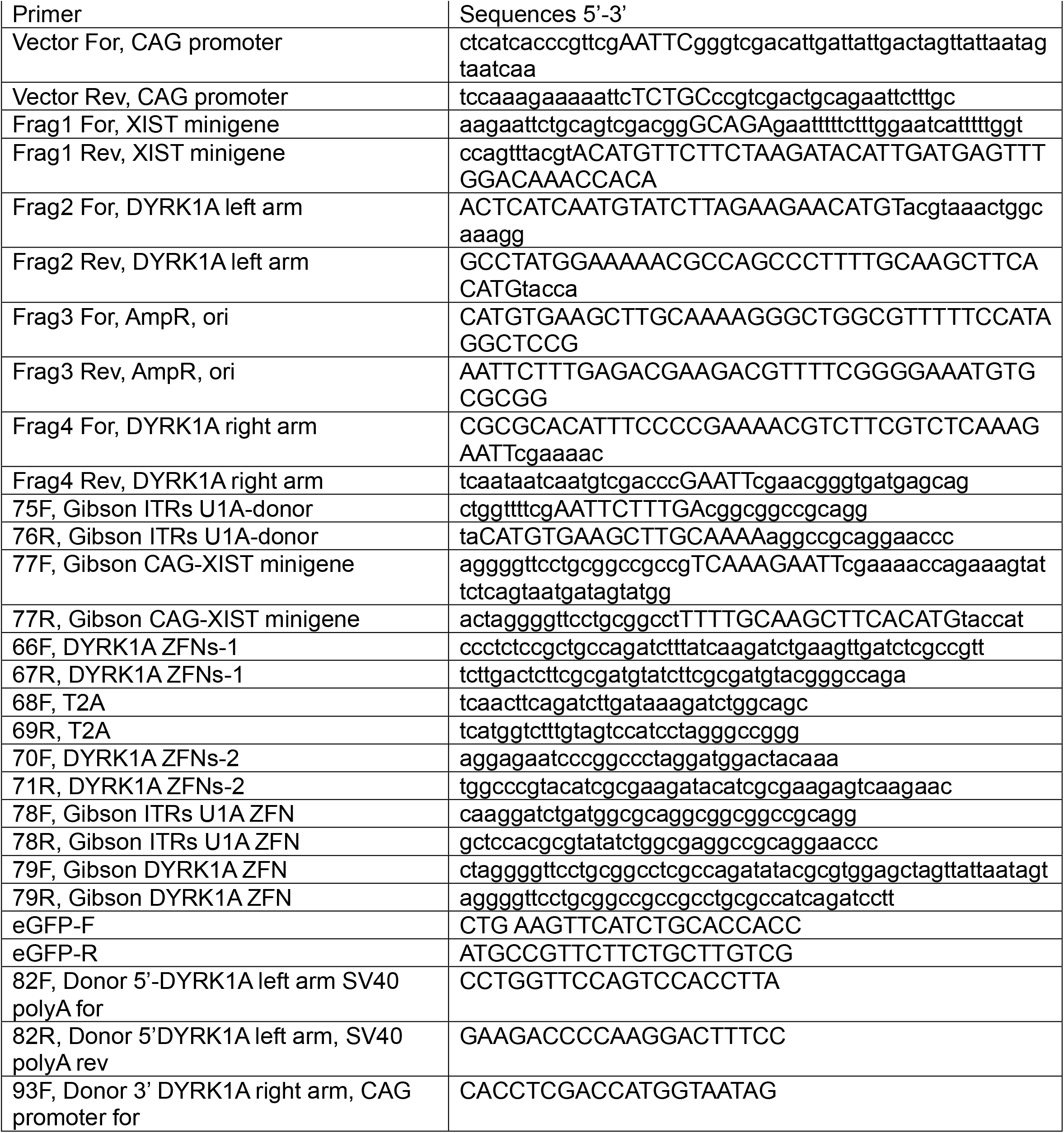
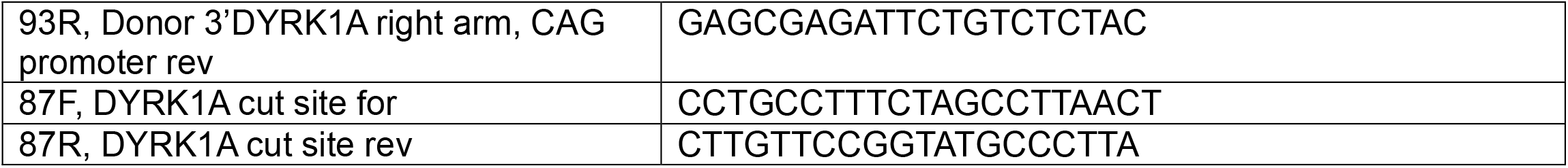

